# Cysteine-Engineered CAR-T Cells to Counter Antigen Escape in B Cell Lymphoma

**DOI:** 10.1101/2025.04.01.646520

**Authors:** Jost Lühle, Simon Krost, Felix Goerdeler, Christian Seitz, Peter H. Seeberger, Oren Moscovitz

**Affiliations:** Department of Biomolecular Systems, Max Planck Institute of Colloids and Interfaces, Potsdam, Germany; Institute of Chemistry and Biochemistry, Freie Universität Berlin, Berlin, Germany; Department of Pediatric Hematology and Oncology, University of Tübingen, Tübingen, Germany; Hopp-Children’s Cancer Center Heidelberg (KiTZ), Heidelberg, Germany; Department of Pediatric Oncology, Hematology, and Immunology, Heidelberg University Hospital, Heidelberg, Germany

**Author notes:** Correspondence should be addressed to Oren Moscovitz (Am Mühlenberg 1, 14476 Potsdam, Germany). Present address: Copenhagen Center for Glycomics, University of Copenhagen, Copenhagen, Denmark (Felix Goerdeler).

## Abstract

Chimeric Antigen Receptor (CAR-) T cell therapy represents a paradigm shift in immunotherapy of hematological cancers. However, selective pressure on cancer cells often leads to suppression of target antigens, eventually causing cancer relapse^1,2^. This so-called antigen escape renders CAR-T cells ineffective, posing a significant clinical challenge^2–5^. Therefore, identifying alternative targets less susceptible to antigen escape is crucial. Here, we describe a novel type of CAR-T cells utilizing cysteine-engineered antibody fragments that target altered redox states on the surface of B cell lymphoma (BCL)^6^. We demonstrate that cysteine-engineered CAR-T cells exhibit specific cytotoxicity *in vitro* against various BCL subtypes, including antigen escape models. Additionally, we show that cysteine engineering, achieved through single amino acid substitution in the state-of-the-art anti-CD19-CAR, enables co-targeting of both CD19-positive and -negative BCL. Our findings introduce a novel class of bifunctional CAR-T cells that target conventional antigens and altered redox states simultaneously, potentially reducing the risk of antigen escape. Abnormal redox states occur in several cancers, including breast and leukemia^7–12^, indicating a broad therapeutic scope.

## Introduction

Immunotherapy represents a key frontier in the development of novel cancer therapeutics. Antibodies and antibody fragments play a central role in immunotherapy as they allow specific targeting of malignant cells. The treatment of hematological cancers has been revolutionized in the last decades by Chimeric Antigen Receptor (CAR-) T cells where the patient’s own T lymphocytes are genetically engineered *ex vivo* with a CAR composed of an extracellular cancer-specific antibody fragment and intracellular signaling domains for triggering T cell activation and cytotoxicity^13^. These CAR-T cells are re-transfused and exert their anticancer activity *in vivo*. To date, there are six different CAR-T cell therapies in clinical use against relapsed or refractory B cell lymphoma (BCL), acute lymphoblastic leukemia (ALL), and multiple myeloma (MM)^14–19^. Except for MM therapies, all approved formats employ CD19, a transmembrane glycoprotein highly expressed by most B cell malignancies. Moreover, the expression of CD19 on healthy cells is restricted to B cells, minimizing on-target/off-tumor effects^20^.

Despite the remarkable advances of CAR-T cell therapy for hematological cancers, the technology still faces several challenges. These include adverse effects like neurotoxicity or cytokine release syndrome, as well as emerging resistance of the cancer1,21,22. The latter is frequently mediated by a mechanism referred to as antigen escape, which is one of the main drawbacks of conventional CAR-T cell therapies. Since they target a specific tumor-associated antigen (TAA), selective pressure upon exposure to CAR-T cells can lead to downregulation or complete loss of the TAA^23,24^. For CD19-targeted CAR-T cells, antigen-negative relapse rates as high as 25% in ALL and BCL have been reported in the clinic^2–5^. Therefore, additional targets for CAR-T cell therapies in hematological cancers are needed. Current attempts focus on alternative targets such as CD20 and bispecific receptors or adapter CAR-T cells to respond dynamically to antigen escape^25–27^. However, these approaches still target specific TAAs, leaving the window open for cancer adaptation and treatment-induced resistance.

An alternative strategy for cancer immunotherapy is to co-target the cancer microenvironment, thereby expanding the target scope and minimizing the risk of evasion mechanisms^28,29^. A common feature of several cancer types, including BCL, is a reduced cellular microenvironment caused by differential expression of disulfide-active enzymes, such as protein disulfide isomerase (PDI) or the thioredoxin system^6–8,11,12,30,31^. These alterations can contribute substantially to disease progression. For example, upregulation of PDI is associated with cancer growth and drug resistance^9,10^, and reduced thiol groups on breast cancer cells facilitate attachment to the endothelium, thereby promoting metastasis^7^. These effects render the reduced cell surface microenvironment an attractive target for therapy. Small molecule PDI inhibitors have been reported to prevent metastasis in breast cancer and prolong the survival of mice transplanted with multiple myeloma^7,32^. On the other hand, increased levels of reduced surface thiols can be exploited for drug delivery, as they can be targeted directly by disulfide-active compounds or antibodies^6,33^.

Previously, we developed a single-domain antibody (nanobody) derived from an alpaca heavy chain-only IgG, which binds several subtypes of BCL^6^. This nanobody, denoted CB2, interacts with altered redox states of the cell surface microenvironment via the non-canonical cysteine 105 in the complementarity-determining region (CDR) 3. Subsequent thiol-mediated uptake of CB2 enables its use as an intracellular drug carrier, but a substantial fraction of the nanobody remains on the cell surface, suggesting its suitability for immunotherapy^6^. Here, we report the development of CAR-T cells with CB2 as antibody fragment, thus directly targeting the cell surface microenvironment of BCL instead of a single antigen. We show that CB2-CARs alone can trigger T cell activation and cytotoxicity against different subtypes of BCL, while healthy human Peripheral Blood Mononuclear Cells (PBMCs) remain unharmed. Furthermore, we demonstrate CB2-CAR-T cell activity against antigen escape models resistant to conventional CAR-T therapies. Next, we performed cysteine engineering on a second, GFP-binding single-domain antibody, highlighting the universal applicability of the method for targeting BCL cells through their abnormal cell surface redox states. Finally, we apply our method to state-of-the-art CD19-CAR-T cells, effectively targeting both CD19-positive and -negative BCL *in vitro* and establish defined criteria for optimization of cysteine-engineered bifunctional CAR constructs.

## Methods

### Design of CAR constructs

The second-generation CAR construct comprising human elongation factor-1 alpha promotor (EF-1α) → CD8 signal peptide (SP) → CB2 domain → human IgG1-Fc hinge → CD28 transmembrane domain (TM) → CD28 costimulatory domain (CD) → CD3ζ signaling domain (SD) → Woodchuck Hepatitis Virus Posttranscriptional Regulatory Element (WPRE) → human cytomegalovirus enhancer and promotor (CMV) → green fluorescent protein (GFP), abbreviated “CB2-CD28-CAR_GFP”, was ordered as synthetic gene in the lentiviral transfer plasmid pLV from VectorBuilder Inc. (Chicago, IL). A GFP control construct (“GFP”) was cloned in-lab via restriction of the plasmid with SpeI followed by self-ligation.

A shorter version of the construct, abbreviated “CB2-CD28-CAR”, was generated via deletion of GFP reporter. Briefly, the complete CAR construct was amplified via polymerase chain reaction (PCR) using a reverse primer with an overhang to insert a 3’ KpnI restriction site. Then, the amplicon and the original backbone were restricted using XbaI (restriction site upstream CB2 domain) and KpnI (restriction site downstream GFP) and ligated. The ^C105S^CB2 mutant of this construct was generated via overlap extension PCR as described by Heckman et al.^34^, leading to “^C105S^CB2-CD28-CAR”. Sequences of the anti-GFP nanobody LaG-16 and its mutant ^S106C^LaG-16 were ordered as synthetic gene fragments from IDT Inc. (Coralville, IA). The sequence was published by Fridy et al.^35^. The gene fragments were inserted into the CD28-CAR backbone via NEBuilder® HiFi DNA Assembly (NEB, Ipswich, MA), leading to “LaG-16-CD28-CAR” and “^S106C^LaG-16-CD28-CAR”.

Furthermore, a second-generation CAR construct comprising granulocyte-macrophage colony-stimulating factor (GM-CSF) SP instead of CD8-SP, CD8 hinge instead of IgG1-Fc hinge, CD8-TM instead of CD28-TM, and 4-1BB-CD instead of CD28-CD was generated. The original plasmid was provided by the Seitz lab, university of Tübingen, with anti-human CD19 single-chain variable fragment FMC63 as extracellular antibody fragment (abbreviated “CD19-CAR”). To clone the construct “CB2-41BB-CAR”, FMC63 was replaced by CB2 via NEBuilder® HiFi DNA Assembly.

The FMC63 cysteine mutants N92C, D156C, S190C, G228C, and S229C were ordered as synthetic DNA fragments (IDT, Coralville, IA, USA) and inserted into the pLV:CD19-CAR backbone via restriction and ligation cloning using EcoRV and NdeI (NEB, Ipswich, MA, USA) according to the manufacturer’s instructions.

All lentiviral constructs used in this study are summarized in Supplementary Fig. 1.

### Lentiviral packaging

HEK293T cells were transfected with pLV (carrying the respective CAR construct), psPAX2 (plasmid was a gift from Didier Trono, Addgene plasmid # 12260; http://n2t.net/addgene:12260; RRID:Addgene_12260), and pMD2.G (plasmid was a gift from Didier Trono, Addgene plasmid # 12259; http://n2t.net/addgene:12259; RRID:Addgene_12259) in a T75 flask with 6.3 µg DNA per plasmid and 30 µL TransIT 293T transfection reagent (Mirus Bio LLC, Madison, WI). Cells were incubated at 37°C for 48 h. Afterwards, viral supernatant was centrifuged at 1500 g for 10 min and filtered through a 0.45 µm filter. For concentration, supernatant was layered on a sucrose buffer (50 mM Tris-HCl, 100 mM NaCl, 0.5 mM EDTA, 10% w/v sucrose, pH 7.4) at 4:1 v/v ratio in a 50 mL tube and centrifuged at 10 000 g/4°C for 4 h. Afterwards, the supernatant was removed, and the viral pellet was resuspended in 500 µL sterile PBS. For storage, virus suspensions were aliquoted and frozen directly at −80°C. Lentivirus concentration protocol was adapted from Jiang et al.^36^.

### CAR-T cell generation

Human whole blood samples were obtained from the German Red Cross (DRK-Blutspendedienst Nord-Ost, Institut Berlin). PBMCs were isolated using SepMate™-50 tubes (Stemcell Technologies, Vancouver, Canada). Afterwards, T cells were isolated using Pan T Cell Isolation Kit (Miltenyi Biotec, Bergisch Gladbach, Germany) and seeded at 1.0 million cells/mL in T cell medium (RPMI 1640, 10% fetal calf serum (FCS), 2 mM L-glutamine, 1 mM pyruvate, 100 U/mL Penicillin-Streptomycin) supplemented with 30 U/mL human Interleukin-2 (IL-2; Peprotech Inc., Cranbury, NJ). 25 µL Dynabeads™ Human T-Activator CD3/CD28 beads (ThermoFisher Scientific, Waltham, MA) per million of T cells were added. One day later, concentrated lentivirus was added in a 1:15 dilution alongside with 5 µg/mL polybrene. On day 2, medium was replaced with fresh T cell medium + IL-2. Cells were monitored daily and maintained between 0.5 and 1.0 million cells/mL. T cells transduced with CB2-CD28-CAR_GFP and GFP control were enriched via fluorescence-activated cell sorting (see respective section) on day 6. All other constructs were not sorted but transduction efficiencies were determined via flow cytometry. Briefly, all nanobody-based CAR-T cells were incubated with MonoRab™ Rabbit Anti-Camelid VHH Cocktail iFluor 647 (1:500; GenScript, Piscataway, NJ) and CD19-CAR-T cells were incubated with Recombinant Human CD19 Fc Chimera Alexa Fluor® 647 (1:500; Bio-Techne, Minneapolis, MN) and/or anti-idiotypic FMC63 scFv Mouse antibody, FITC conjugate (1:50; ACROBiosystems, Newark, DE) for FMC63 mutants at room temperature (RT) for 30 min. Samples were measured on a FACSCanto™ II Flow Cytometry System (BD, Franklin Lakes, NJ). 24 h before experiments, T cells were transferred into medium without IL-2. For long-term storage, CAR-T cells were cryo-preserved in liquid nitrogen with 90% FCS, 10% dimethyl sulfoxide as freezing medium.

### Fluorescence-Activated Cell Sorting

T cells transduced with CB2-CD28-CAR_GFP were stained with MonoRab™ Rabbit Anti-Camelid VHH Cocktail iFluor 647 at 4°C for 30 min. GFP control T cells were resuspended in PBS. Subsequently, cells were sorted for nanobody-positive or GFP-positive cells respectively, using a FACSMelody™ Cell Sorter (BD, Franklin Lakes, NJ). Sorting efficiency was determined afterwards using the same staining pattern (Supplementary Fig. 2).

### Apoptosis assays

Non-transduced, sorted GFP control and sorted CB2-CAR-T cells were co-cultured with SC-1 cells in different Effector to Target (E:T) ratios (1:1, 3:1, 5:1 and 9:1). Each condition was prepared in triplicates. SC-1 cell numbers were constant (5×10^3^ cells/well). Co-cultivation took place in a 96-well plate for 72 h at 37°C. For analysis, cells were stained with anti-CD3-APC antibody (1:200; BioLegend, San Diego, CA) to distinguish B cell lymphoma (BCL) and T cells, Annexin V-PE (1:100; BioLegend, San Diego, CA) to stain early apoptotic cells and 7-AAD Viability Staining Solution (1:100; ThermoFisher Scientific, Waltham, MA) to stain late apoptotic cells. Incubation took place in Annexin V Binding Buffer (BioLegend, San Diego, CA) at RT for 15 min. Cells from each well were analyzed separately on a FACSCanto™ II. Target cells were gated for single cells and SC-1 cells (CD3-/GFP-events; Supplementary Fig. 4). Average frequencies of early and late apoptotic SC-1 cells were compared between the different conditions and E:T ratios.

Apoptosis assays with healthy human PBMCs were performed as described above, but PBMCs were stained with CellTrace™ Far Red Cell Proliferation Kit (ThermoFisher Scientific, Waltham, MA) prior to co-culture to distinguish them from CAR-T cells and replace the use of anti-CD3 antibody. Therefore, healthy PBMCs were gated for CellTrace™+/GFP-events.

Significance of differences was assessed via 2-way ANOVA followed by Tukey’s post-hoc test.

### Cytokine secretion assays

Cytokine levels were measured via enzyme-linked immunosorbent assay (ELISA). Supernatants (9:1 ratio) of the co-culture assays described above were collected and levels of IFN-γ and TNF-α were determined in triplicates using Human IFN-γ and Human TNF-α Mini TMB ELISA Kits (Peprotech Inc., Cranbury, NJ) or IFN gamma Human ELISA Kit (ThermoFisher Scientific, Waltham, MA), according to the manufacturer’s instructions. Samples of T cells only were included to determine baseline cytokine secretion. Significance of differences was assessed via 2-way ANOVA followed by Tukey’s post-hoc test.

### T cell activation marker expression

T cells were co-cultivated with SC-1 cells in an E:T ratio of 9:1 in triplicates as described above. Subsequently, each replicate was split into three equal fractions and stained with one of the antibodies for marker detection (PE/Cyanine7 anti-human CD25, PE/Cyanine7 anti-human CD107a (LAMP-1) or PE/Cyanine7 anti-human CD69; all BioLegend, San Diego, CA). Anti-CD3-APC antibody (BioLegend, San Diego, CA) was included in all samples to gate for T cells. Baselines of marker expression were determined with T cells only samples. Marker expression was measured on a FACSCanto™ II. Expression levels of T cells co-cultivated with SC-1 cells were subtracted with expression levels of T cells only. Significance of differences for each marker was assessed via 1-way ANOVA followed by Tukey’s post-hoc test.

### Cell culture

SC-1 and HEK293T cell lines were purchased from DSMZ, Braunschweig, Germany. JeKo-1 and its knockouts were kindly provided by the Seitz lab, university of Tübingen. All cell lines were maintained in a humidified incubator at 37°C and 5% CO2. HEK293T cells were cultured in DMEM medium supplemented with 10% FCS, 2 mM L-glutamine, and 100 U/mL Penicillin-Streptomycin. All BCL lines were cultured in RPMI 1640 medium supplemented with 10% FCS, 2 mM L-glutamine, 1 mM pyruvate and 100 U/mL Penicillin-Streptomycin. For culture conditions of CAR-T cells, see respective section. All cell culture media and supplements were purchased from PAN-Biotech GmbH, Aidenbach, Germany. Cell cultures were tested negative for mycoplasma via PCR monthly.

### JeKo-1 lymphoma knockout cell lines

JeKo-1 cells and the knockout lines JeKo-1 CD19-KO and JeKo-1 CD20-KO, all expressing firefly luciferase, were generated by the Seitz lab, university of Tübingen, as described elsewhere^27^. Knockout efficiencies were determined via flow cytometry (Supplementary Fig. 5) incubating with PE anti-human CD19 Antibody (1:100) and APC anti-human CD20 Antibody (1:100; both BioLegend, San Diego, CA) at RT for 1 h.

### Luciferase-expressing SC-1 cells

SC-1 cells expressing firefly luciferase and GFP were generated in-lab via lentiviral transduction. Lentiviral particles were produced as described above with SIN40C.SFFV.Luciferase.IRES.GFP as transfer plasmid (plasmid was a gift from Jan-Henning Klusmann, Addgene plasmid # 169308; http://n2t.net/addgene:169308; RRID:Addgene_169308)^37^. One million SC-1 cells were transduced with 120 µL concentrated lentivirus and 8 µg/mL polybrene. On day 8, cells were single-cell sorted for GFP-positive cells using a FACSMelody™. Expanding clones were screened on day 23 for uniform GFP expression. One clone was selected, further expanded and cryo-preserved. The SC-1 origin was confirmed via flow cytometry with PE anti-human CD19 Antibody (1:100; BioLegend, San Diego, CA), confirming uniform CD19 and GFP expression of the clone (Supplementary Fig. 6).

### Luciferase-based cytotoxicity assays

Non-transduced and CAR-T cells were co-cultivated with BCL in different E:T ratios (1:1, 3:1, 5:1 and 9:1). Each condition was prepared in triplicates. BCL cell numbers were constant (5×10^3^ cells/well). BCL only samples were included for lysis control. Differences in CAR-T cell transduction efficiencies were considered by adjusting T cell numbers to reach the same amounts of CAR-positive cells among all constructs. Co-cultivation took place in a 96-well plate at 37°C for 24 h (SC-1) or 48 h (all other BCL). For analysis, 100 µg/mL D-luciferin potassium salt (Abcam, Cambridge, UK) was added to each well. For lysis control wells, 1% NP-40 was added. Cells were incubated at 37°C for 1 h. Afterwards, relative light units (RLU) were measured in a CLARIOstar Plus plate reader (BMG LABTECH GmbH, Ortenberg, Germany) with an exposure time of 10 seconds per well. Normalized specific lysis was calculated by normalizing test RLUs to target cells co-cultured with non-transduced T cells (NT) at the respective E:T ratio (background cytotoxicity) and lysis control wells (maximal killing) using the following equation:

Normalized spec. lysis [%] = 100 × [(NT RLU – test RLU) / (NT RLU – lysis control RLU)]

Data from independent experiments was pooled by calculating the grand mean and pooled standard error. Significance of differences was assessed via 2-way ANOVA followed by Tukey’s post-hoc test.

### Protein structure analysis

The crystal structure of FMC63 in complex with CD19 was previously published^38^, and can be retrieved from the protein database PDB (Entry ID 7URV). The structure was modified and analyzed using the software PyMOL. One parameter that was assessed is the relative solvent-accessible surface area (SASA), a theoretical value indicating the relative surface area of each amino acid that is in contact with the surrounding solvent^39^. For the calculation of SASA values, the structure of FMC63 was displayed without CD19. SASA values were computed via the command “PyMOL>get_sasa_relative polymer”. To calculate the minimal distance to the antigen, the measurement wizard tool in PyMOL was used to determine the distance (in Ångström [Å]) between the respective amino acid’s side chain and the closest atom of CD19.

### Recombinant expression of nanobodies

Histidine-tagged LaG-16 expression plasmid “pET21-pelB-LaG-16” was a gift from Michael Rout (Addgene plasmid #172746; http://n2t.net/addgene:172746; RRID:Addgene_172746)^35^. The S106C mutation was introduced by overlap-extension PCR as described elsewhere^34^. Both plasmids were transformed into ArcticExpress (DE3) *Escherichia coli* cells (Agilent Technologies, Santa Clara, CA), and protein expression was induced in lysogenic broth medium with 1 mM isopropyl-β-D-1-thiogalactopyranoside at 12°C overnight. Bacteria were lysed with 5 freeze-thaw cycles, alternating between a 37°C water bath and dry ice. Nanobody was then purified in two steps, first by Nickel-NTA agarose resin (ThermoFisher Scientific, Waltham, MA) followed by size exclusion chromatography on an ÄKTApurifier using an S200 column (Cytiva, Marlborough, MA). All steps and purity of the final product were validated by sodium dodecyl sulfate poly-acrylamide gel electrophoresis (SDS-PAGE, Supplementary Fig. 7). CB2 and ^C105S^CB2 were expressed and purified as described elsewhere^6^.

### Flow cytometry nanobody binding assays

One million SC-1 cells per sample were incubated with 24 µM nanobody solution or PBS (negative control) at RT for 1 h. Afterwards, all samples were incubated with MonoRab™ Rabbit Anti-Camelid VHH Cocktail iFluor 647 at RT for 1 h. Samples were measured on a FACSCanto™ II.

### Ethics statements

Blood sampling from healthy human donors and downstream experiments were approved by the ethics council of the Max Planck Society (Application number 2021_08).

## Results

### CB2-CAR-T cells specifically target BCL

We demonstrated previously that CB2 nanobodies specifically target BCL cells via thiol interactions^6^. Therefore, we hypothesized that CB2 is also suitable to guide CAR-T cells towards BCL in a thiol-dependent manner (Fig. 1a). We designed a second-generation CAR construct with CB2 as antigen fragment, human IgG1-Fc as hinge, and CD28 as transmembrane and costimulatory domains. GFP was part of the same vector and served as a reporter protein (CB2-CD28-CAR_GFP, Fig. 1c). We successfully activated and transduced primary human T cells with lentivirus carrying the CB2-CD28-CAR_GFP or GFP control construct and enriched CAR-T cells via fluorescence-activated cell sorting (Fig. 1b, Supplementary Fig. 2).

**Fig. 1.**
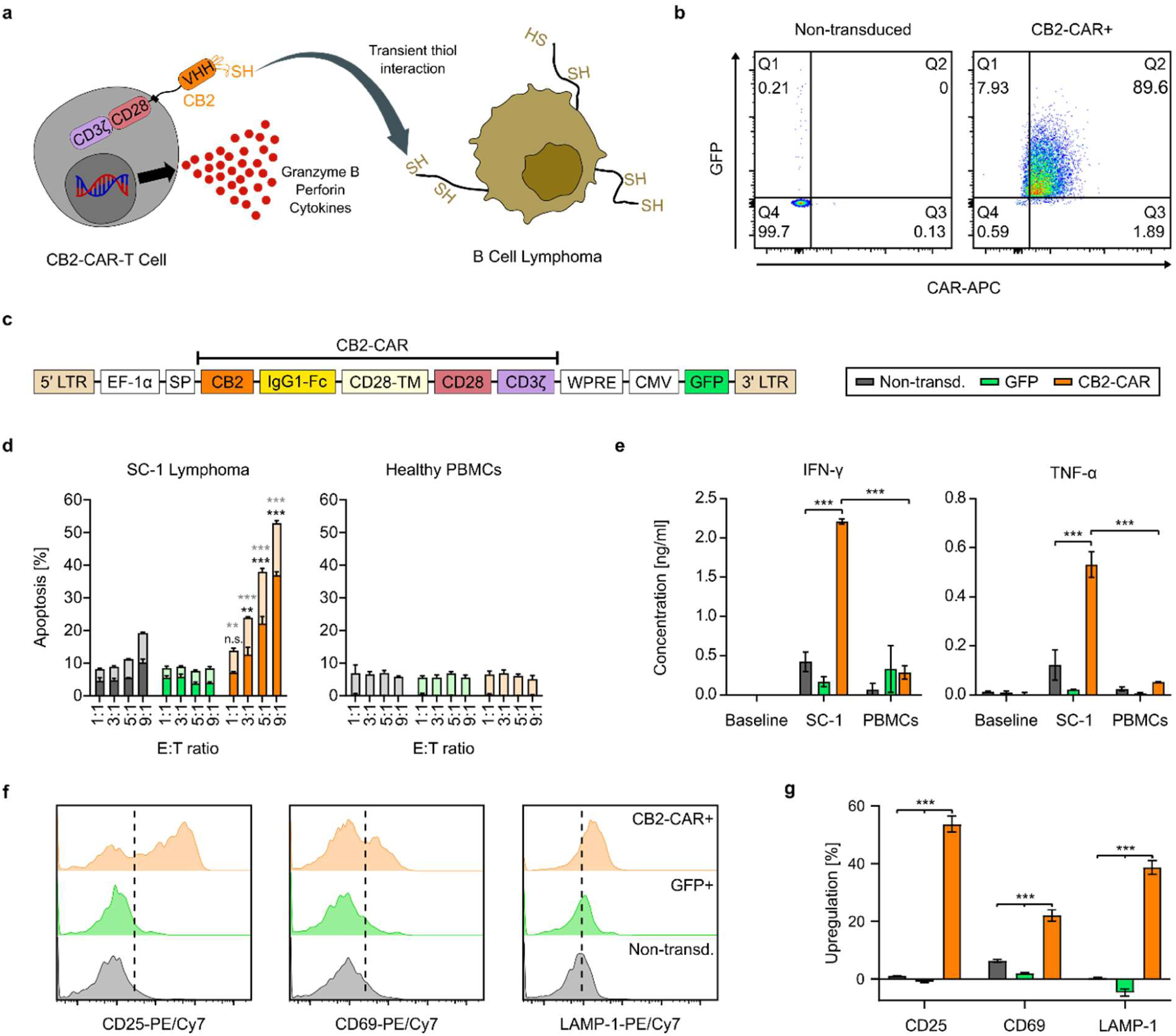
CB2-CAR-T cells exhibit cytotoxicity against B cell lymphoma and undergo T cell activation. **a**, Schematic representation of thiol-dependent CB2-CAR-T cell activity against BCL. **b**, Transduction efficiency of sorted CB2-CAR-T cells on day 6 after transduction, determined via flow cytometry. Representative plots are shown. Percentages of the individual populations are indicated in the graphs. **c**, Domain structure of CB2-CD28-CAR_GFP construct. LTR – long terminal repeat, EF-1α – human elongation factor-1 alpha promotor, SP – signal peptide, IgG1-Fc – human IgG1-Fc hinge, CD28-TM – CD28 transmembrane domain, CD28 – CD28 costimulatory domain, CD3ζ – CD3ζ signaling domain, WPRE – Woodchuck Hepatitis Virus Posttranscriptional Regulatory Element, CMV – human cytomegalovirus enhancer and promotor, GFP – green fluorescent protein. **d**, Flow cytometry-based apoptosis assays. Stacked bars represent the different types of apoptosis (bottom part for late apoptosis and top part for early apoptosis). Mean values ± SEM of n=3 replicates are shown. Significance was assessed via 2-way ANOVA followed by Tukey’s post-hoc test. Bottom significance levels refer to differences in late apoptosis and top significance levels to differences in early apoptosis compared to both controls. **e**, Cytokine secretion assays (ELISA). Baselines represent T cells cultured without target cells under the same conditions. Mean values ± SEM of n=3 replicates are shown, and significance of differences was assessed via 2-way ANOVA followed by Tukey’s post-hoc test. **f**, Representative flow cytometry plots for the analysis of T cell activation marker expression. **g**, Quantification of T cell activation marker expression. Fractions of positive cells shown in (**f**) were quantified by subtracting the baseline expression of each marker. Mean values ± SEM of n=3 replicates are shown, and significance of differences was assessed via 1-way ANOVA followed by Tukey’s post-hoc test for each marker individually. Significance levels in all subfigures: n.s. – not significant; * – p<0.05; ** – p<0.01; *** – p<0.001.

To test CAR-T cell activity, we co-cultured them with SC-1 cells (follicular lymphoma) that have an abnormal cell surface redox status^6^ at different effector to target (E:T) ratios and measured apoptosis of the target cells via flow cytometry. CB2-CAR-T cells exhibited significant concentration-dependent cytotoxicity compared with non-transduced and GFP control T cells (>50% at 9:1 ratio). In contrast, co-cultures with healthy human PBMCs showed no cytotoxic activity (Fig. 1d). To further profile CAR-T cell activation, we performed enzyme-linked immunosorbent assays (ELISAs) to test the secretion of pro-inflammatory cytokines IFN-γ and TNF-α upon co-incubation with SC-1 cells. In agreement with the apoptosis assay results, CB2-CAR-T cells secreted approximately four-fold higher amounts of both cytokines than control T cells. In parallel, healthy PBMCs did not trigger cytokine secretion (Fig. 1e). To confirm activity on the effector cell level, we assessed upregulation of T cell activation markers, namely CD25 and CD69, and the degranulation marker LAMP-1 via flow cytometry. All three markers were significantly upregulated in the case of CB2-CAR-T cells co-incubated with SC-1 (54%, 22%, and 39% for CD25, CD69, and LAMP-1, respectively), confirming T cell activation and degranulation (Fig. 1f, g).

These results demonstrate CB2-CAR-T cell activation upon co-culture with SC-1 lymphoma. More precisely, CAR-T cells exhibit cytotoxicity, secrete pro-inflammatory cytokines, and upregulate T cell activation markers, suggesting that CB2-CAR-T cells target BCL via an antigen-independent mechanism solely based on redox interactions.

### Cysteine engineering redirects CARs towards BCL

We chose to engineer the non-cancer-related anti-GFP nanobody LaG-16^35^ with a cysteine in CDR3 as a proof-of-concept for the antigen-independent mechanism. The cysteine-to-serine mutant ^C105S^CB2 does not bind to SC-1 cells in flow cytometry^6^. Reversely, we introduced a cysteine at position 106 in the sequence of LaG-16 (^S106C^LaG-16) and probed SC-1 cell binding via flow cytometry. Indeed, ^S106C^LaG-16 showed the same binding pattern as wild-type CB2, whereas wild-type LaG-16, like ^C105S^CB2, did not bind SC-1 cells (Fig. 2a, b). Encouraged by this result, we hypothesized that the activity of CB2-CAR-T cells depends solely on the exposed cysteine in CDR3, and that CARs of different specificity can be redirected towards BCL when endowed with such a cysteine.

**Fig. 2.**
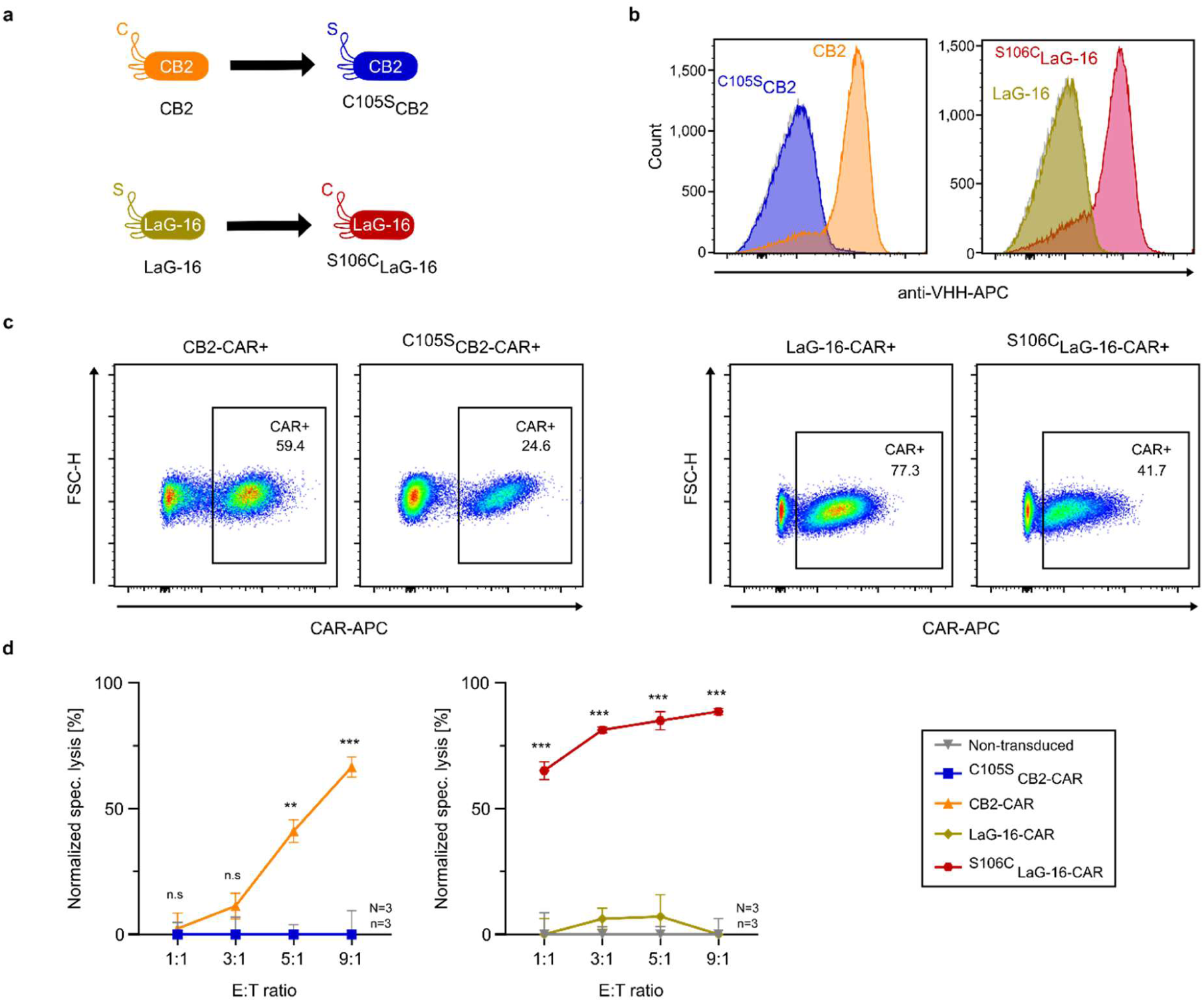
Cysteine-engineered nanobodies redirect CAR-T cells to target B cell lymphoma. **a**, Illustration of investigated nanobodies and corresponding mutations. **b**, Flow cytometry assays to test binding of nanobodies to SC-1 cells. Grey histograms in the background correspond to secondary antibody-only controls. Representative plots of n=3 experiments are shown. **c**, Determination of transduction efficiencies of CB2-CAR- and LaG-16-CAR-T cells on day 6 after transduction via flow cytometry. Representative plots are shown. Percentages of CAR+ T cells are indicated in the graphs. **d**, Luciferase-based cytotoxicity assays of CAR-T cells co-cultured with SC-1 lymphoma at different E:T ratios for 24 hours. Grand mean values ± pooled SEM of N=3, n=3 replicates are shown. Tested constructs are non-transduced (▾), ^C105S^CB2-CAR (▪), CB2-CAR (▴), LaG-16-CAR (⧫), and ^S106C^Lag-16-CAR (⬣). Significance of differences was assessed via 2-way ANOVA followed by Tukey’s post-hoc test (n.s. – not significant; * – p<0.05; ** – p<0.01; *** – p<0.001).

To test this hypothesis, we generated CAR-T cells expressing wild-type CB2, ^C105S^CB2, wild-type LaG-16, and ^S106C^LaG-16 (Supplementary Fig. 1). Transduction efficiencies differed repeatedly between constructs (Fig. 2c), but cell numbers were adjusted in all assays to reach equivalent amounts of effector cells. We performed luciferase-based cytotoxicity assays with SC-1 cells to compare the activity of the different constructs. In accordance with the cell binding assays, both constructs harboring a cysteine in CDR3 (CB2-CAR and ^S106C^LaG-16-CAR) exhibited significant cytotoxicity (>50% at 9:1 ratio). In contrast, the serine-containing counterparts (^C105S^CB2-CAR and LaG-16-CAR) did not show any activity. Interestingly, the activity of ^S106C^LaG-16-CAR-T cells was higher at each E:T ratio compared to CB2-CAR-T cells (Fig. 2d).

These results demonstrate how non-cancer-related nanobody-based CAR-T cells can be redirected to target BCL by engineering the CDR3 with a single cysteine, whereas replacing this cysteine with serine leads to a complete loss of activity. Additionally, the superior performance of ^S106C^LaG-16 as a CAR compared to CB2 reveals intriguing differences in functionality that are influenced by the position of cysteine in the nanobody sequence. Based on these findings, we inspected the therapeutic scope of cysteine-engineered CAR-T cells against other BCL subtypes and antigen escape models.

### Cysteine-dependent targeting of antigen escape models

To compare the activity of CB2-CAR-T cells with a standard CD19-CAR-T therapy, we changed the receptor configuration to match the clinically used Kymriah® format (CD19-CAR, Fig. 3a, b), which we used as a control. More specifically, we introduced CD8-derived hinge and transmembrane domains and the 4-1BB-derived costimulatory domain (CB2-CAR, Fig. 3b) and generated CB2-CAR-T and CD19-CAR-T cells by lentiviral transduction with high efficiencies (>60%, Fig. 3c). To evaluate cytotoxicity, we co-cultured CAR-T cells with SC-1 cells, JeKo-1 cells (mantle cell lymphoma), and JeKo-1 cells with knocked-out CD19 or CD20 antigens to model antigen escape (Supplementary Fig. 5) and assessed the supernatants for IFN-γ secretion via ELISA. As negative controls, we included healthy PBMCs and red blood cells (RBCs). In accordance with our previous results, CB2-CAR-T cells showed a significant increase in IFN-γ secretion against all cancer cell lines. Notably, IFN-γ levels against CD19-negative cells reached approximately 3 ng/mL, while there was no secretion from CD19-CAR-T cells (Fig. 3d). Healthy PBMCs did not trigger any IFN-γ secretion of CB2-CAR-T cells. In contrast, CD19-CAR-T cells showed a small but significant increase in IFN-γ levels, presumably against healthy B cells (Fig. 3d).

**Fig. 3.**
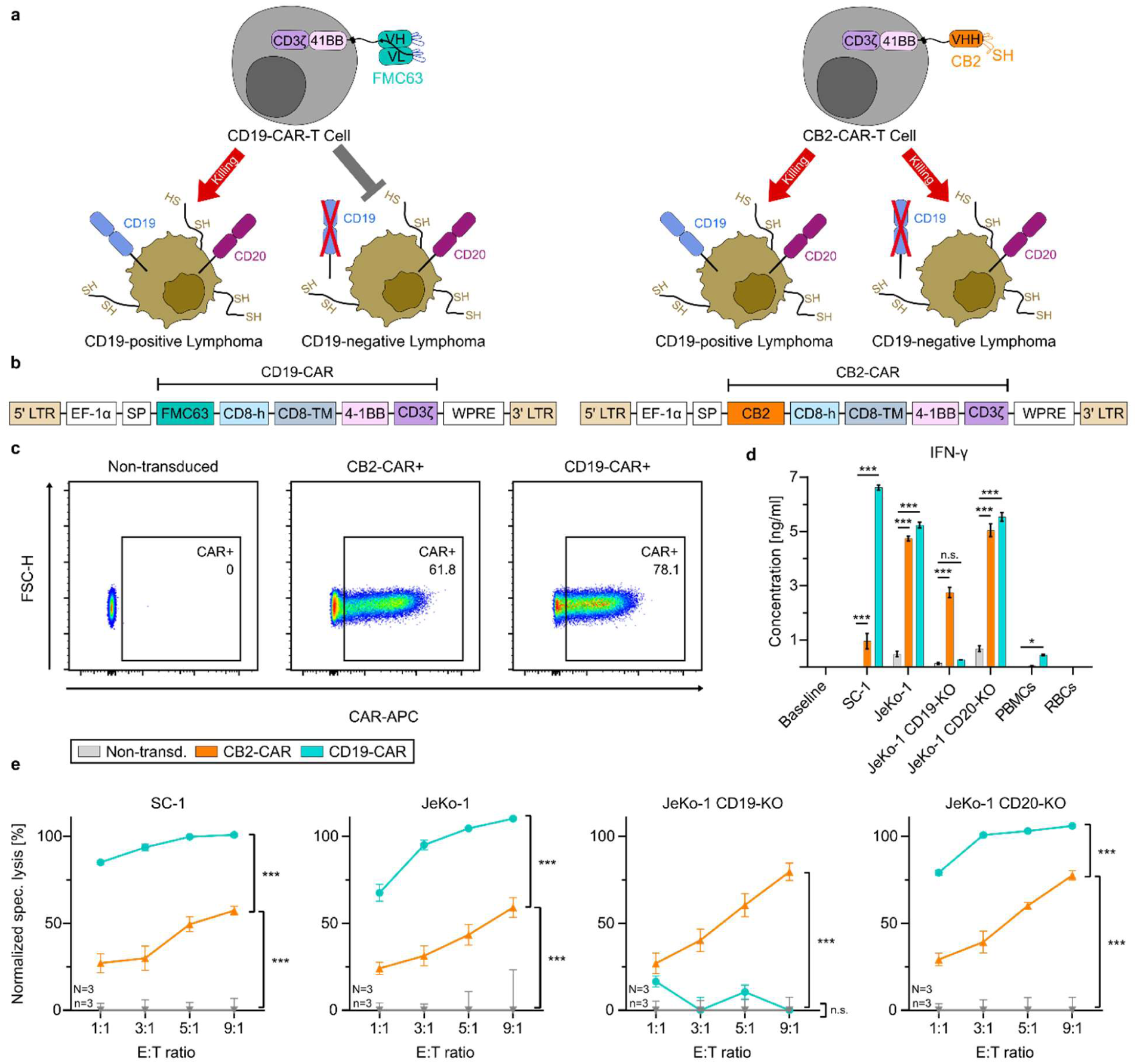
CB2-CAR-T cells efficiently target B cell lymphoma subtypes and antigen escape models. **a**, Schematic illustration of CD19-independent targeting by CB2-CAR-T cells. **b**, Domain structures of CB2-CAR and CD19-CAR. LTR – long terminal repeat, EF-1α – human elongation factor-1 alpha promotor, SP – signal peptide, CD8-h – CD8 hinge, CD8-TM – CD8 transmembrane domain, 4-1BB – 4-1BB costimulatory domain, CD3ζ – CD3ζ signaling domain, WPRE – Woodchuck Hepatitis Virus Posttranscriptional Regulatory Element. **c**, Determination of transduction efficiencies of CB2-CAR and CD19-CAR on day 6 after transduction via flow cytometry. Representative plots are shown. Percentages of CAR+ T cells are indicated in the graphs. **d**, IFN-γ secretion assay (ELISA). Baseline represents T cells cultured without target cells under the same conditions. Mean values ± SEM of n=3 replicates are shown, and significance of differences was assessed via 2-way ANOVA followed by Tukey’s post-hoc test. **e**, Luciferase-based cytotoxicity assays of different lymphoma cell lines co-cultured with CAR-T cells at different E:T ratios for 24 hours (SC-1) or 48 hours (JeKo-1, JeKo-1 CD19-KO, and JeKo-1 CD20-KO). Grand mean values ± pooled SEM of N=3, n=3 replicates are shown. Tested constructs are non-transduced (▾), CB2-CAR (▴), and CD19-CAR (●). Significance of differences was assessed via 2-way ANOVA followed by Tukey’s post-hoc test. Significance levels in all subfigures: n.s. – not significant; * – p<0.05; ** – p<0.01; *** – p<0.001.

After confirming IFN-γ secretion, we aimed to quantify the cytotoxicity of CB2-CAR-T cells against different BCL subtypes and antigen escape models via luciferase assays. To demonstrate clinical relevance, we pooled data from three independent blood donors for CAR-T cells (N=3, n=3). CB2-CAR-T cells exhibited significant cytotoxicity in a concentration-dependent manner against all tested BCL subtypes (Fig. 3e). Furthermore, JeKo-1 CD19-knockout cells were targeted with high activity (≈80% specific lysis at 9:1 ratio), while remaining resistant to CD19-CAR-T cells. For other BCL lines, the cytotoxicity of CB2-CAR-T cells was lower compared to CD19-CAR-T cells but consistently reached values above 50% (Fig. 3e).

In conclusion, CB2-CAR-T cells are active against different BCL subtypes as well as antigen escape models and a BCL line resistant to conventional CD19-directed CAR-T cells can be targeted efficiently. Therefore, we investigated whether cysteine engineering could be applied to the clinically used scFv-based CD19-CAR-T cells, enabling them to simultaneously target both CD19 and cancer-associated redox states.

### Bifunctionality of Cysteine-engineered CD19-CAR-T cells

To assess whether cysteine engineering can be applied to scFv-based CAR-T cells, we selected five candidate residues in the sequence of CD19-specific FMC63 for individual cysteine replacement. The residues were chosen according to their location in the different CDRs, and their different biochemical properties. For this, we included two serine residues (S190 and S229), a glycine (G228), an asparagine (N92), and an aspartic acid (D156). N92 is located in light chain (LC) CDR3, D156 in heavy chain (HC) CDR1, S190 in HC-CDR2, and G228 and S229 in HC-CDR3 (Fig. 4a, b).

**Fig. 4.**
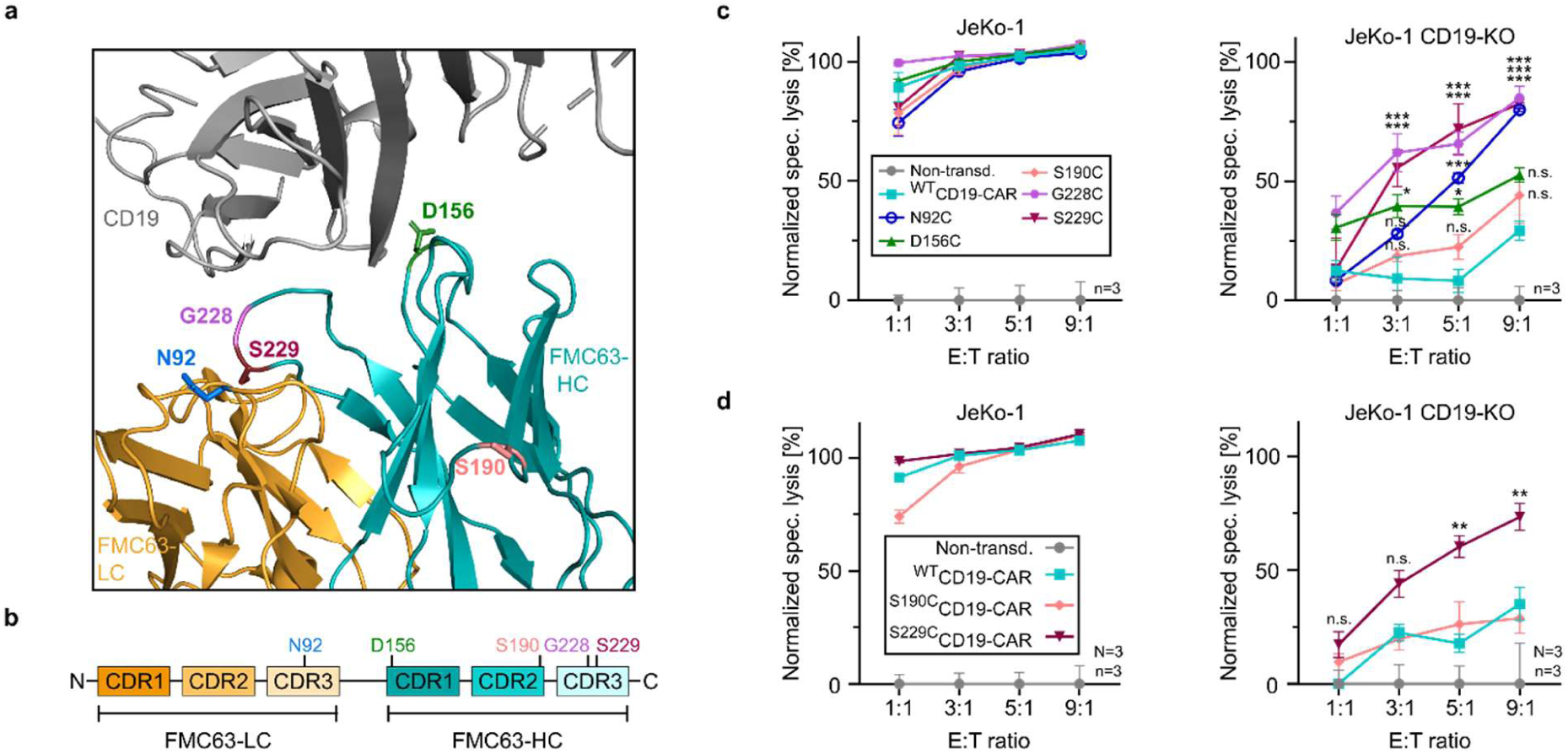
Bifunctional activity of cysteine-engineered CD19-CAR-T cells. **a**, Location of the mutated residues N92, D156, S190, G228, and S229 within the three-dimensional structure of FMC63 in complex with CD19 (data retrieved from PDB entry ID 7URV). **b**, Schematic representation of the individual mutated residues within the linear sequence of FMC63. LC – light chain, HC – heavy chain. **c,d**, Luciferase-based cytotoxicity assays of CD19-CAR-T cells co-cultured with JeKo-1 lymphoma or JeKo-1 CD19-knockout at different E:T ratios for 48 hours. Tested constructs are non-transduced (●), ^WT^CD19-CAR (▪), ^N92C^CD19-CAR (○), ^D156C^CD19-CAR (▴), S190/^C^CD19-CAR (♦), ^G228C^CD19-CAR (⬣), and ^S229C^CD19-CAR (▼). Mean values ± SEM of n=3 (**c**) or N=3, n=3 (**d**) replicates are shown. Significance of differences was assessed via 2-way ANOVA followed by Tukey’s post-hoc test (n.s. – not significant; * – p<0.05; ** – p<0.01; *** – p<0.001). All indicated significance levels are in comparison to ^WT^CD19-CAR-T cells.

We generated mutated CD19-CAR-T cells and determined transduction efficiencies using an anti-idiotypic antibody. Efficiencies >50% were achieved for all mutants besides G228C (Supplementary Fig. 3a). However, incubation of ^G228C^CD19-CAR-T cells with recombinant, fluorescently labeled CD19 resulted in a detectable fluorescence shift, indicating presence of the CAR (Supplementary Fig. 3b). We performed co-cultures of all mutated CAR-T cells, as well as wild-type CD19-CAR-T cells, with JeKo-1 and JeKo-1 CD19-KO cells, respectively. All tested constructs exhibited a similar cytotoxic activity against CD19-positive JeKo-1 cells, indicating CAR expression as well as retained target affinity of all mutants (Fig. 4c, left panel). Moreover, most cysteine-engineered CARs effectively targeted CD19-negative lymphoma, in contrast to the neglectable activity of ^WT^CD19-CAR-T cells. Interestingly, two constructs with adjacent cysteine mutations (G228C and S229C) showed similar cytotoxicity (Fig. 4c, right panel). Their comparable activity, despite the differences in the substituted amino acid, suggests that the position of the mutation has a higher impact than the chemical properties of the replaced residue. This notion was underscored as only one of the serine mutants (S229C) showed a significant gain in cysteine-dependent activity, while the other (S190C), located in a different CDR, did not (Fig. 4c, right panel).

Based on these results, we sought to identify parameters to elucidate the variations in activity observed among cysteine-engineered CARs. As the position of the mutation within a CDR ensures accessibility of the introduced cysteine, we restricted candidate residues to the CDRs. Additionally, we propose two structural parameters that can serve as a basis for rational cysteine engineering, based on a published FMC63-CD19 structure^38^. The first parameter is the relative solvent-accessible surface area (SASA), a theoretical value indicating the relative surface area of each amino acid in contact with the surrounding solvent. We computed relative SASA values for all mutated residues (summarized in Supplementary Table 1). When comparing SASA values to the specific lysis at the highest E:T ratio, mutations with a lower SASA (<0.5) show higher cytotoxicity, indicating that less solvent-exposed residues are advantageous. Furthermore, we suggest the minimal distance to the antigen as another structural parameter for selecting candidate residues. We compared these distance values (in Ångström [Å]) with the specific lysis at the highest E:T ratio (Supplementary Table 1). Our experimental data show that distances of 4-8 Å achieved a maximum killing of ≥80%, with diminished but existing activity of the aspartic acid (D156C) mutant (5.7 Å). In contrast, the S190C mutant, 16.7 Å away from the antigen, showed no significant activity (Supplementary Table 1, Fig. 4c). These results suggest prioritizing residues closer to the antigen binding site as a selection criterion.

We chose the serine-to-cysteine mutant S229C for subsequent experiments, as it showed high cytotoxicity against JeKo-1 CD19-KO cells (Fig. 4c) and can be detected more reliably than the similarly active G228C mutant (Supplementary Fig. 3a). To show clinical relevance, we repeated the co-cultures with ^WT^CD19-CAR-T and ^S229C^CD19-CAR-T cells generated from three independent blood donors (N=3, n=3). In accordance with previous results, ^S229C^CD19-CAR-T cells showed significant cytotoxic activity, also against CD19-negative lymphoma (Fig. 4d). In contrast, ^S190C^CD19-CAR-T cells did not exhibit an increased cytotoxicity against knockout target cells, serving as negative control (Fig. 4d).

In conclusion, we successfully applied cysteine engineering to the clinically used scFv-based CD19-CAR-T cells, demonstrating that substituting a single amino acid with cysteine enables these cells to target both antigen-positive and antigen-negative lymphoma. This introduces a novel structure-to-function approach for developing bifunctional CAR-T cells that simultaneously target a specific antigen and abnormal surface redox states of cancer cells.

## Discussion

CAR-T cell therapies have shown great success in the treatment of hematological cancers, including B cell lymphoma. Nevertheless, the ability of cancers to dynamically adapt to changes in the environment leads to antigen escape as one of the main reasons for treatment failure^2,24^. Here, we describe a novel type of antigen-independent CAR-T cells that exhibit anticancer activity by interacting directly with the cellular microenvironment of BCL, thereby potentially reducing the risk of antigen escape.

By employing the cysteine-exposing nanobody CB2, we successfully produced CAR-T cells that specifically target BCL via thiol interactions. CB2-CAR-T cell activity was confirmed by cytotoxicity assays, secretion of pro-inflammatory cytokines, and activation marker expression (Fig. 1). We demonstrate how thiol interactions via cysteine-105 of CB2 are sufficient to trigger T cell activation and cytotoxicity, and result in an activation profile that is comparable to conventional protein-targeted CAR-T cells^40,41^. Moreover, we show that healthy human PBMCs do not trigger CB2-CAR-T cell activation, indicating specificity towards BCL (Fig. 1d, e). These results imply that BCL’s altered cell surface redox states are sufficient to serve as a novel target for CAR-T cells.

The interesting CB2-CAR-T cell activity led to investigation into whether a cysteine-engineered CDR3 can confer activity against BCL to any nanobody-based CAR, employing the anti-GFP nanobody LaG-16 as a proof-of-concept. Cell binding assays to SC-1 cells demonstrated that introducing a single cysteine in CDR3 causes binding to BCL, while replacement by serine abolishes it (Fig. 2a, b). Corroborating this notion, cysteine-modified CAR-T cells exhibit significant, concentration-dependent cytotoxicity, while serine counterparts are completely inactive (Fig. 2d). These experiments confirm that cysteine engineering of a non-cancer-related nanobody can redirect T cells to target BCL when expressed as part of a CAR. Furthermore, the inactivity of serine-exposing counterparts supports a model where CAR-T cell activation relies exclusively on a single cysteine in CDR3.

The proof-of-concept experiments with LaG-16 point to many potential applications of cysteine-engineered, bispecific antibodies as independent antibody modules, part of bispecific T cell engagers (BiTEs), or as CARs guiding T cells. Since activity depends on a single amino acid exchange, basically any TAA-specific antibody or antibody fragment can be modified to recognize both the original antigen and altered redox states. In this way, producing novel bifunctional antibodies is conceivable. Since the cysteine modification is located in CDR3 and is part of the paratope, the position must be chosen carefully to minimize interference with antigen binding. The overall higher activity of ^S106C^LaG-16-CAR-T compared to CB2-CAR-T cells (Fig. 2d) further suggests a careful design of the exact cysteine position for optimal CAR-T cell activity.

Next, we evaluated the therapeutic scope of our CB2-CAR-T cells and compared it to the clinical state-of-the-art anti-CD19 CAR-T cell therapy Kymriah®. Indeed, CB2-CAR-T cells displayed significant activity against different BCL lines, including JeKo-1 cells with knocked-out CD19 and CD20 (Fig. 3e). Remarkably, CD19-knockout cells were targeted efficiently by CB2-CAR-T cells, while remaining resistant to CD19-CAR-T cells. These results emphasize that cysteine-engineered CAR-T cells are promising candidates for countering antigen escape in BCL, as they remain cytotoxic against antigen escape models and activity does not depend on a single surface antigen. To support these findings, IFN-γ ELISAs were performed with co-culture supernatants. While there was significant IFN-γ secretion of CB2-CAR-T cells against all types of lymphoma, including CD19-knockouts, no secretion was measured against healthy PBMCs and RBCs (Fig. 3d). As conventional CD19-directed CAR-T cell therapies also affect healthy B cells^42^, these results suggest that by using CAR-T cells solely acting through an engineered cysteine, on-target/off-tumor effects are potentially reduced.

Encouraged by our results, we speculated that the cysteine engineering approach can be applied to the clinically used CAR-T cell therapy Kymriah®. As these CD19-directed CAR-T cells are based on the scFv FMC63 that has two domains in contrast to a nanobody, we systematically tested five candidate cysteine mutations to establish the optimal position within the three-dimensional structure. We identified several mutations that target CD19-expressing lymphoma with efficiencies similar to ^WT^CD19-CAR-T cells while demonstrating a parallel activity against CD19-negative lymphoma (Fig. 4c, d). Evaluating different mutations provided valuable insights into structural parameters that serve as a basis for application of cysteine engineering to other CARs. The distance to the antigen binding site appears essential, as a more distant mutation (S190C) shows a diminished activity (Supplementary Table 1, Fig. 4c), presumably due to impaired signal transduction. Residues with an increased relative solvent-accessible surface area (SASA) were less active against CD19-negative lymphoma (Supplementary Table 1, Fig. 4c), supposedly due to increased spontaneous oxidation, as reported for cysteines with higher SASA values^43^. We introduce a type of engineered CD19-CAR-T cells, targeting both their specific antigen as well as altered cell surface redox states of BCL. This mechanism has the potential to improve existing therapies by reducing the risk of antigen escape through the cysteine-mediated elimination of antigen-negative cancer subpopulations. Furthermore, we suggest two structural parameters to guide the development of cysteine-modified CAR-T cells against additional antigens and cancer types. Whether cysteine-engineered CAR-T cells are effective *in vivo* remains to be determined and will be subject of further research. However, several studies have successfully targeted altered cancer redox states *in vivo* without significant systemic effects, underscoring the therapeutic potential of this approach^33,44–46^.

While this study focuses on B cell malignancies, the therapeutic scope of cysteine-engineered CAR-T cells is broader, as altered redox states of the cell surface microenvironment have also been reported for other cancer types. Breast cancer, for example, has been shown to upregulate PDI and expose increased levels of surface thiols^7,30^, as we also demonstrated by showing specific binding of CB2 to breast cancer cell lines^6^. It remains to be explored whether targeting altered redox states in solid tumors offers advantages compared to conventional CAR-T cells. This is particularly relevant when targeting potential antigens that exhibit inconsistent and varying expression levels in patients, such as the disialoganglioside GD2 antigen in triple-negative breast cancer patients^47,48^. Nevertheless, our current study suggests that using CB2 or other cysteine-engineered CAR-T cells for breast cancer treatment is conceivable.

In conclusion, our findings present a novel CAR-T cell therapy approach, demonstrating effective targeting of B cell lymphoma by engaging the cellular redox microenvironment through cysteine-exposing CARs. This suggests a potential bypass for antigen escape, opening additional avenues for the development of bifunctional single-chain and single-domain antibodies with transformative implications for various malignancies.

## Acknowledgements

Financial support by the Max Planck Society is gratefully acknowledged. This work was further supported by Deutsche Forschungsgemeinschaft (RTG2046 for J.L. and F.G).

We thank Prof. Dr. Helge Ewers for his help and support.

## Author contributions

J.L. and O.M. designed the concept. J.L. established the methods, planned and performed the experiments, and wrote the manuscript with contributions from all other authors. F.G. performed the mutation and binding assays of recombinant LaG-16. C.S. and S.K. produced and provided the JeKo-1 reporter cell line, the JeKo-1 CD19- and CD20-knockout lines, provided the lentiviral transfer plasmid for CD19-CAR, and supported this work with experimental advice. O.M. conceived the original idea. O.M. and P.H.S. supervised the project.

