## Supplementary Information for "Cysteine-Engineered CAR-T Cells to Counter Antigen Escape in B Cell Lymphoma"

### Supplementary Figures

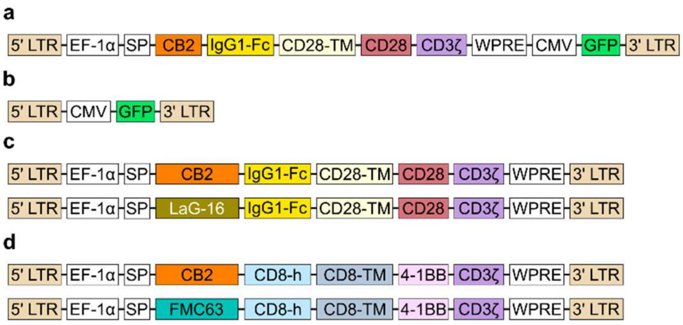


**Supplementary Fig. 1 | Overview of CAR constructs used in this study. a**, CB2-CD28-CAR_GFP construct. **b**, GFP control construct. **c**, CD28-CAR constructs. **d**, 41BB CAR constructs. LTR – long terminal repeat, EF-1α – human elongation factor-1 alpha promotor, SP – signal peptide, IgG1-Fc – human IgG1-Fc hinge, CD28-TM – CD28 transmembrane domain, CD28 – CD28 costimulatory domain, CD3ζ – CD3ζ signaling domain, WPRE – Woodchuck Hepatitis Virus Posttranscriptional Regulatory Element, CMV – human cytomegalovirus enhancer and promotor, GFP – green fluorescent protein, CD8-h – CD8 hinge, CD8-TM – CD8 transmembrane domain, 4 1BB – 4 1BB costimulatory domain.

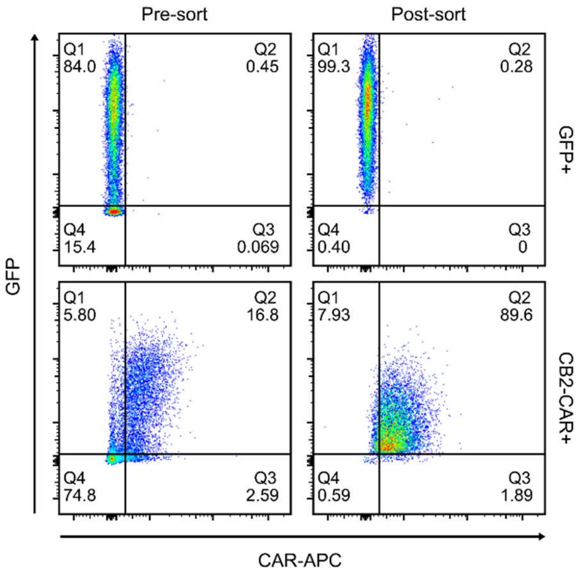


**Supplementary Fig. 2 | Sorting efficiency of CB2-CD28-CAR_GFP and GFP control T cells.** Pre-sorting (left panels) and post-sorting (right panels) efficiencies were measured via flow cytometry. CAR-APC and GFP signals were assessed.

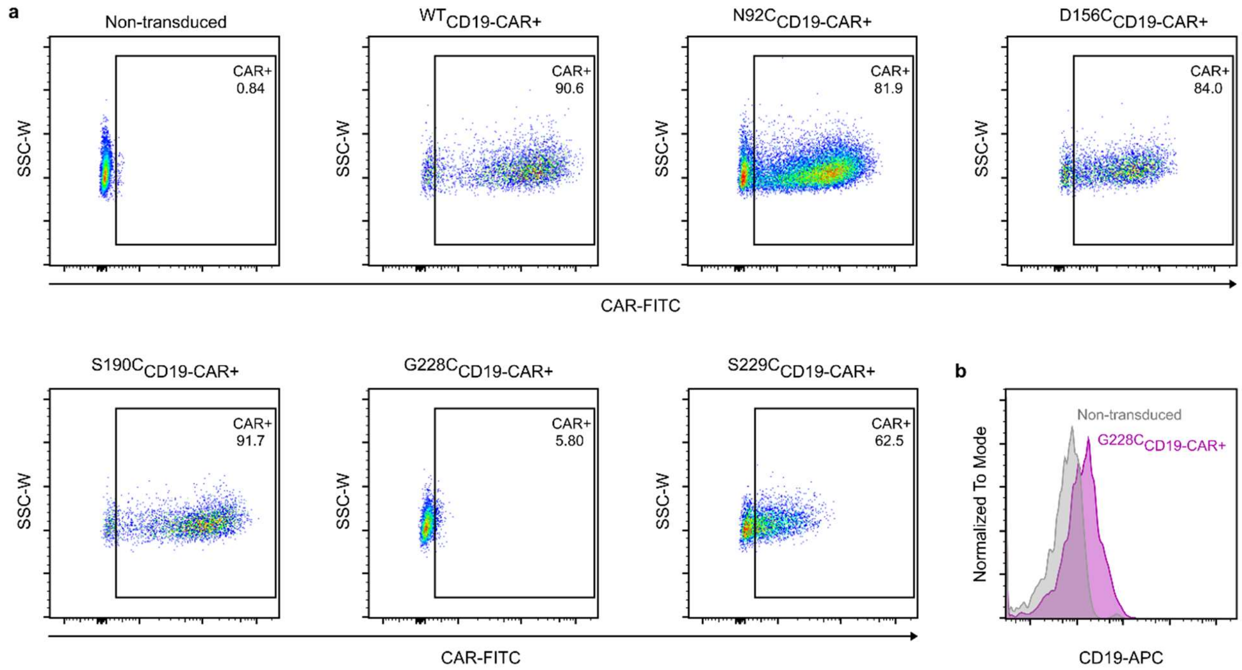


**Supplementary Fig. 3 | Transduction efficiencies of WT and mutated CD19-CAR-T cells.** **a**, Determination of transduction efficiencies of CD19-CAR variants on day 6 after transduction via flow cytometry. Representative plots are shown. Percentages of CAR+ T cells are indicated in the graphs. **b**, Verification of CAR expression on $^{G228C}$CD19-CAR-T cells via staining with fluorescent CD19 (flow cytometry).

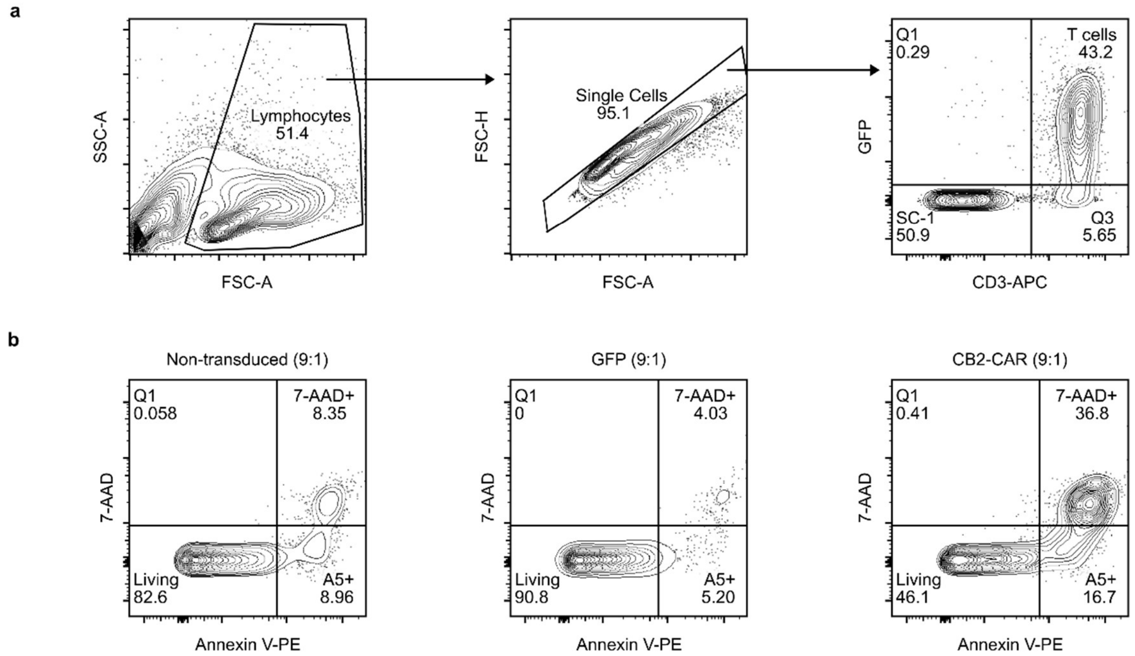


**Supplementary Fig. 4 | Gating strategy for flow cytometry-based apoptosis assays. a**, Gating of single target cells (SC-1). Cells were gated for lymphocytes (left panel), single cells (middle panel), and SC-1 cells (GFP-/CD3+; right panel). **b**, Gating of apoptotic target cells. Single SC-1 cells (gated in A) were further gated for Annexin V-PE+/7-AAD- cells (early apoptotic cells) and Annexin V-PE+/7-AAD+ cells (late apoptotic cells). Example for effector to target ratio of 9:1 is shown. FSC – forward scatter, SSC – side scatter.

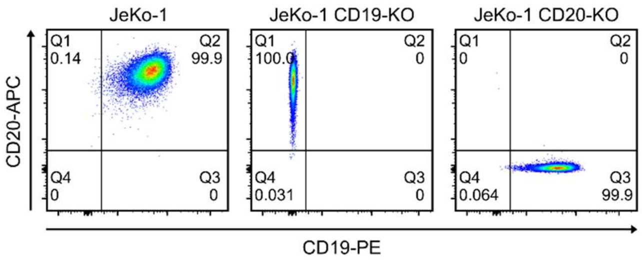


**Supplementary Fig. 5 | Knockout efficiencies of JeKo-1 cell lines.** Efficiencies were determined via flow cytometry. Gates were adjusted according to non-stained control (not shown). Left panel: JeKo-1, middle panel: JeKo-1 CD19-knockout, right panel: JeKo-1 CD20-knockout.

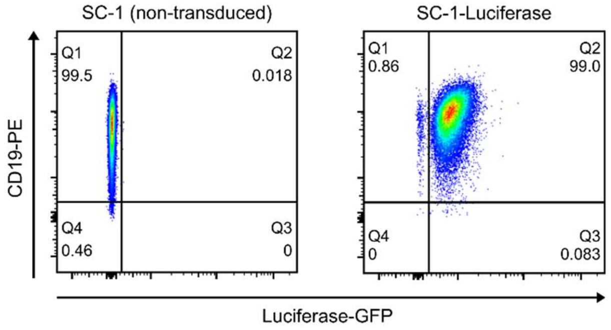


**Supplementary Fig. 6 | Validation of firefly luciferase-expressing SC-1 cells.** SC-1 cell line was transduced with lentivirus carrying firefly luciferase and GFP. CD19 and Luciferase-GFP expression was assessed via flow cytometry to confirm successful generation of luciferase-expressing SC-1 cells.

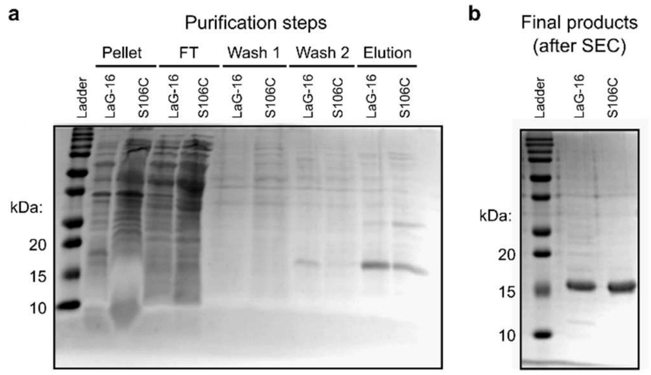


**Supplementary Fig. 7 | Purification of recombinant nanobodies LaG-16 and $^{S106C}$LaG-16 from *Escherichia coli*. a**, Sodium dodecyl sulfate poly-acrylamide gel electrophoresis (SDS-PAGE) of purification steps. Samples from the different steps of Nickel-NTA affinity chromatography are shown. Nanobodies are visible in the gel at a size of approximately 15 kilodalton. **b**, Purity of final products. FT – flow through, kDa – kilodalton, SEC – size exclusion chromatography.

#### Supplementary Tables

**Supplementary Table 1 | Structural parameters of CD19-CAR mutants and corresponding cytotoxic activity**. Specific lysis values refer to co-cultures against JeKo-1 CD19-KO at an effector to target (E:T) ratio of 9:1 (shown in Fig. 4c).

| CD19-CAR mutant | Specific lysis at 9:1 E:T ratio [%] | Relative solvent-accessible surface area | Minimal distance to antigen [Å] |
|---|---|---|---|
| G228C | 85 | 0.30 | 4.2 |
| S229C | 83 | 0.03 | 7.9 |
| N92C | 80 | 0.34 | 5.0 |
| D156C | 53 | 0.82 | 5.7 |
| S190C | 44 | 0.93 | 16.7 |
